# Exon 44 skipping in Duchenne muscular dystrophy: NS-089/NCNP-02, an antisense oligonucleotide with a novel connected-sequence design

**DOI:** 10.1101/2023.02.23.529798

**Authors:** Naoki Watanabe, Yuichiro Tone, Tetsuya Nagata, Satoru Masuda, Takashi Saito, Norio Motohashi, Kazuchika Takagaki, Yoshitsugu Aoki, Shin’ichi Takeda

**Affiliations:** Discovery Research Laboratories in Tsukuba, Nippon Shinyaku Co., Ltd, Tsukuba, Ibaraki, Japan; Department of Molecular Therapy, National Institute of Neuroscience, National Center of Neurology and Psychiatry (NCNP), Kodaira, Tokyo, Japan; Department of Neurology and Neurological Science, Graduate School of Medicine, Tokyo Medical and Dental University, Tokyo, Japan

**Author notes:** Correspondence should be addressed to S.T., National Institute of Neuroscience, 4-1-1 Ogawa-higashi, Kodaira, Tokyo 187-8502, Japan. These authors contributed equally to this work (Naoki Watanabe, Yuichiro Tone, Tetsuya Nagata).

**Keywords:** Duchenne muscular dystrophy, dystrophin, exon 44, exon skipping, antisense therapeutics, morpholino

## Abstract

Exon-skipping therapy mediated by antisense oligonucleotides (ASOs) is expected to provide a therapeutic option for Duchenne muscular dystrophy (DMD). ASOs for exon skipping reported so far target a single continuous sequence in or around the target exon. In the present study, we investigated ASOs for exon 44 skipping (applicable to approximately 6% of all DMD patients) to improve activity by using a novel ASO design incorporating two connected sequences. Phosphorodiamidate morpholino oligomers targeting two separate sequences in exon 44 were created to simultaneously target two splicing regulators in exon 44, and their exon 44 skipping was measured. NS-089/NCNP-02 showed the highest skipping activity among the oligomers. NS-089/NCNP-02 also induced exon 44 skipping and dystrophin protein expression in cells from a DMD patient to whom exon 44 skipping is applicable. We also assessed the *in vivo* activity of NS-089/NCNP-02 by intravenous administration to cynomolgus monkeys. NS-089/NCNP-02 induced exon 44 skipping in skeletal and cardiac muscle of cynomolgus monkeys. In conclusion, NS-089/NCNP-02, an ASO with a novel connected-sequence design, showed both *in vitro* and *in vivo* exon-skipping activity.

## INTRODUCTION

Duchenne muscular dystrophy (DMD) is an X-chromosome-linked progressive hereditary muscle disease caused by mutations in the *DMD* gene, which codes for dystrophin, a protein expressed underneath the muscle-cell membrane. According to estimates from newborn screening,^1^ DMD affects 1 in 3,500 newborn boys and is the most severe and common form of muscular dystrophy. Mutations in the *DMD* gene can lead to either the severe DMD or the milder Becker muscular dystrophy (BMD), depending on whether the translational reading frame is lost or maintained, respectively.^2^ DMD is mainly caused by out-of-frame mutations in the *DMD* gene, which result in a lack of dystrophin. BMD, in contrast, is mostly caused by in-frame mutations, which result in a reduced amount of dystrophin or a reduction in its molecular size.

Exon skipping is a therapeutic approach that uses antisense oligonucleotides (ASOs) to modify pre-mRNA splicing, thereby correcting the translational reading frame and resulting in an internally truncated but partially functional protein. Eteplirsen (brand name, Exondys 51) is the first approved antisense drug for DMD in the USA, and it provides a treatment option for DMD patients who are amenable to exon 51 skipping (approximately 14% of all DMD patients).^3^ Golodirsen (brand name, Vyondys 53), approved in the USA, and viltolarsen (brand name, Viltepso), approved in Japan and the USA, are phosphorodiamidate morpholino oligomers (PMOs) for the treatment of DMD patients who are amenable to exon 53 skipping (approximately 8% of all DMD patients).^4,5^ Casimersen (brand name, Amondys 45), designed to skip exon 45, was approved in the USA.^6^

Exon 44 skipping is mainly applicable to patients with deletions in the *DMD* gene consisting of exons 35–43, 45 or 45–54. Exon 44 skipping is applicable to approximately 6% of DMD patients,^7^ and useful antisense sequences against exon 44 are already known.^8–14^ It has been reported that the exon skipping activity could be improved by targeting two sequences around the target exon using a mixture of two ASOs.^15,16^ However, existing ASOs target only a single continuous sequence in or around exon 44. In this study, we describe the screening of ASOs based on a novel sequence design created by combining two targeting sequences within a single ASO. We report the discovery of NS-089/NCNP-02, a PMO for exon 44 skipping, and present a non-clinical pharmacology study of NS-089/NCNP-02 in human rhabdomyosarcoma (RD) cells, DMD patient-derived cells and cynomolgus monkeys.

## RESULTS

### Screening of ASOs targeting two separate sequences in exon 44

ASOs 20–30 nt in length targeting two separate sequences in exon 44 were investigated. Sequence screening involved three steps as described below (see Figure 1) and was evaluated on the basis of exon 44 skipping activity in RD cells.

**Figure 1.**
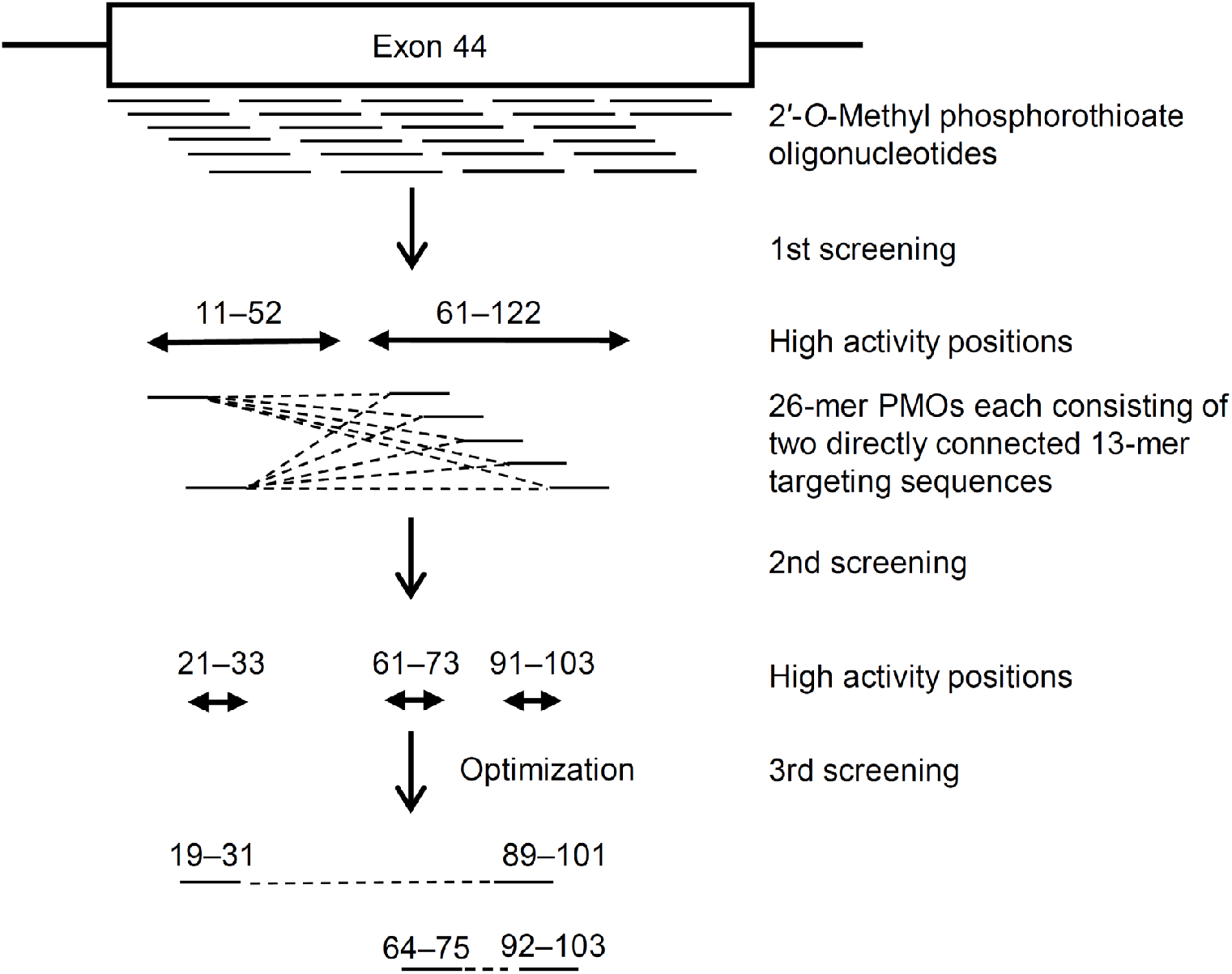
Overview of strategy for screening sequences for exon 44 skipping.

For the first screening, 2′-*O*-methyl phosphorothioate oligonucleotides targeting sequences within exon 44 were synthesized as an overlapping series of twenty-six 22-mer oligomers. These oligomers provided coverage of positions 1 to 147 of this 148-nt exon in 5-nt increments from the 3′ to the 5′ end. The highest exon 44 skipping efficiency was observed for oligomers targeting sequences from positions 11–52 and 61–122 of the exon (data not shown). For the second screening, ten 26-mer PMOs, each consisting of two directly connected 13-mer targeting sequences, one selected from the region of positions 11–33 and the other from the region of positions 61–113 of exon 44, were synthesized, and their exon 44 skipping activity was measured. All 10 PMOs showed skipping activity, and those consisting of the three combinations of two directly connected sequences from the group of sequences corresponding to positions 21–33, 61–73 and 91–103 showed high skipping activity. For the third screening, to optimize the sequence, 18–30-mer PMOs consisting of two connected targeting sequences corresponding to the region of the sequences selected for the second screening were synthesized, and their exon 44 skipping activity was measured. The PMOs all showed high skipping activity. The highest skipping activity was shown by a 26-mer PMO consisting of sequences from positions 19–31 and 89–101 of exon 44, and a 24-mer PMO consisting of sequences from positions 64–75 and 92–103 of exon 44. Since the activities of the 26-mer and the 24-mer were similar, the 24-mer, NS-089/NCNP-02, was selected for further study. The exon skipping efficiency of NS-089/NCNP-02 measured in RD cells showed a dose-dependent increase with an EC_50_ value of 0.83 μmol/L (95% confidence interval, 0.71–0.98 μmol/L; Figure 2A).

**Figure 2.**
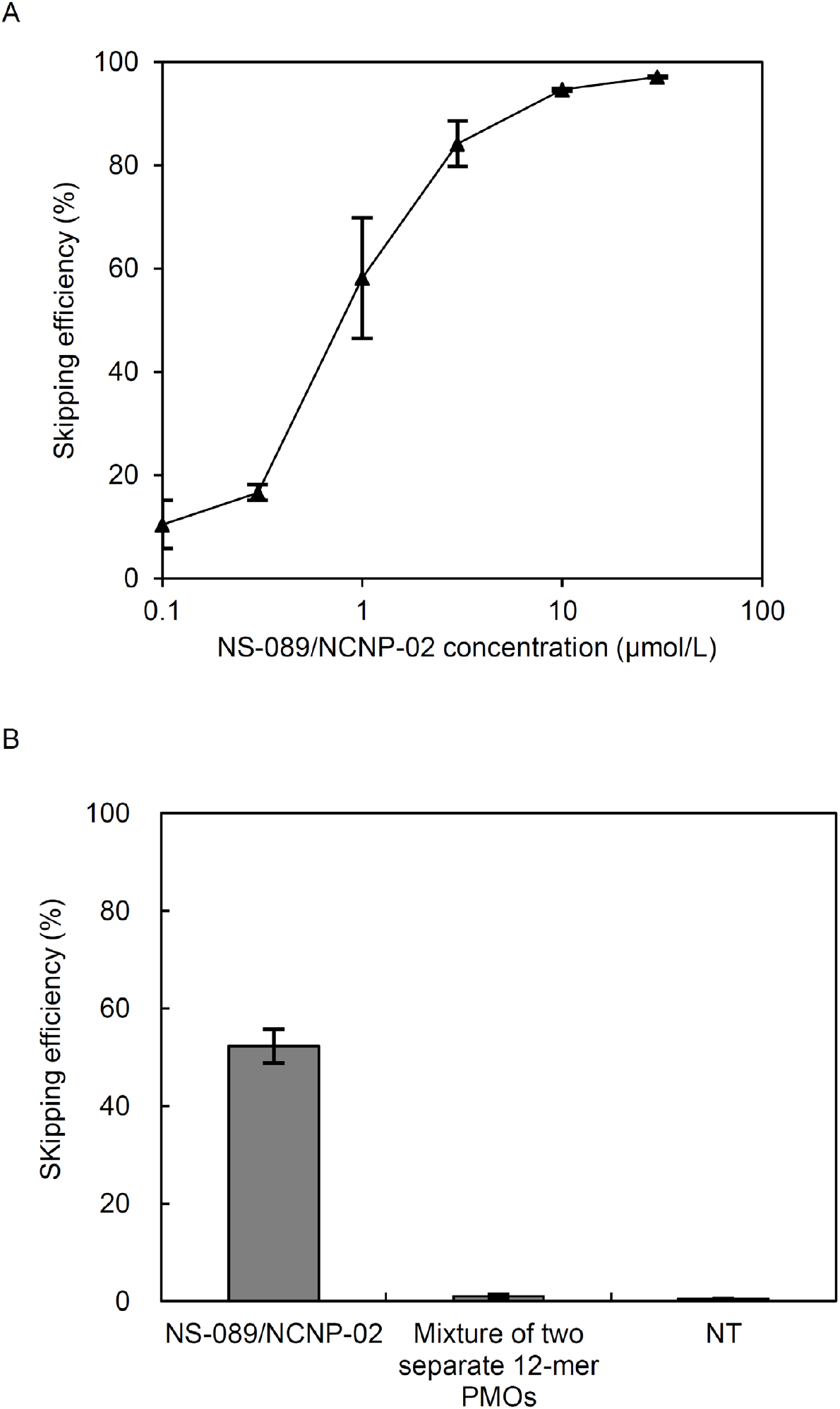
Exon 44 skipping activity of NS-089/NCNP-02 and a mixture of its partial sequences. (A) NS-089/NCNP-02 at concentrations of 0.1, 0.3, 1, 3, 10 and 30 μmol/L or (B) NS-089/NCNP-02 or a mixture of its two partial sequences was transfected into RD cells with Nucleofector, and exon skipping was measured after three days using RT-PCR. NT, non-treated cells used as a negative control. Each point and bar shows the mean ± standard deviation (n = 3).

To investigate whether the high activity of NS-089/NCNP-02 is related to the fact that the targeting sequences are part of the same molecule, its exon 44 skipping activity in RD cells was compared to that of a mixture of two separate 12-mer PMOs corresponding to the same sequences (positions 64–75 and 92–103 of exon 44). NS-089/NCNP-02 showed a skipping efficiency of 52.3% at a concentration of 1 μmol/L, whereas the mixture of two separate 12-mer PMOs each at a concentration of 1 μmol/L showed little activity, about the same as in non-treated cells (Figure 2B).

### Exon 44 skipping activity of NS-089/NCNP-02 in cells derived from patients with DMD

We next measured the exon 44 skipping activity of NS-089/NCNP-02 in cells derived from a DMD patient with deletion of exon 45. Patient fibroblasts were transduced with the human *MYOD* gene to induce differentiation into myotubes. The myotubes were then treated with NS-089/NCNP-02 at final concentrations of 0.01, 0.03, 0.1, 0.3, 1, 3 and 10 μmol/L for two days, after which the differentiation medium was replaced with medium containing no NS-089/NCNP-02. The skipping activity was measured by RT-PCR seven days after the beginning of treatment (Figure 3A). NS-089/NCNP-02 induced exon 44 skipping in the cells with an EC_50_ value of 0.33 μmol/L (95% confidence interval, 0.22–0.51 μmol/L; Figure 3B).

**Figure 3.**
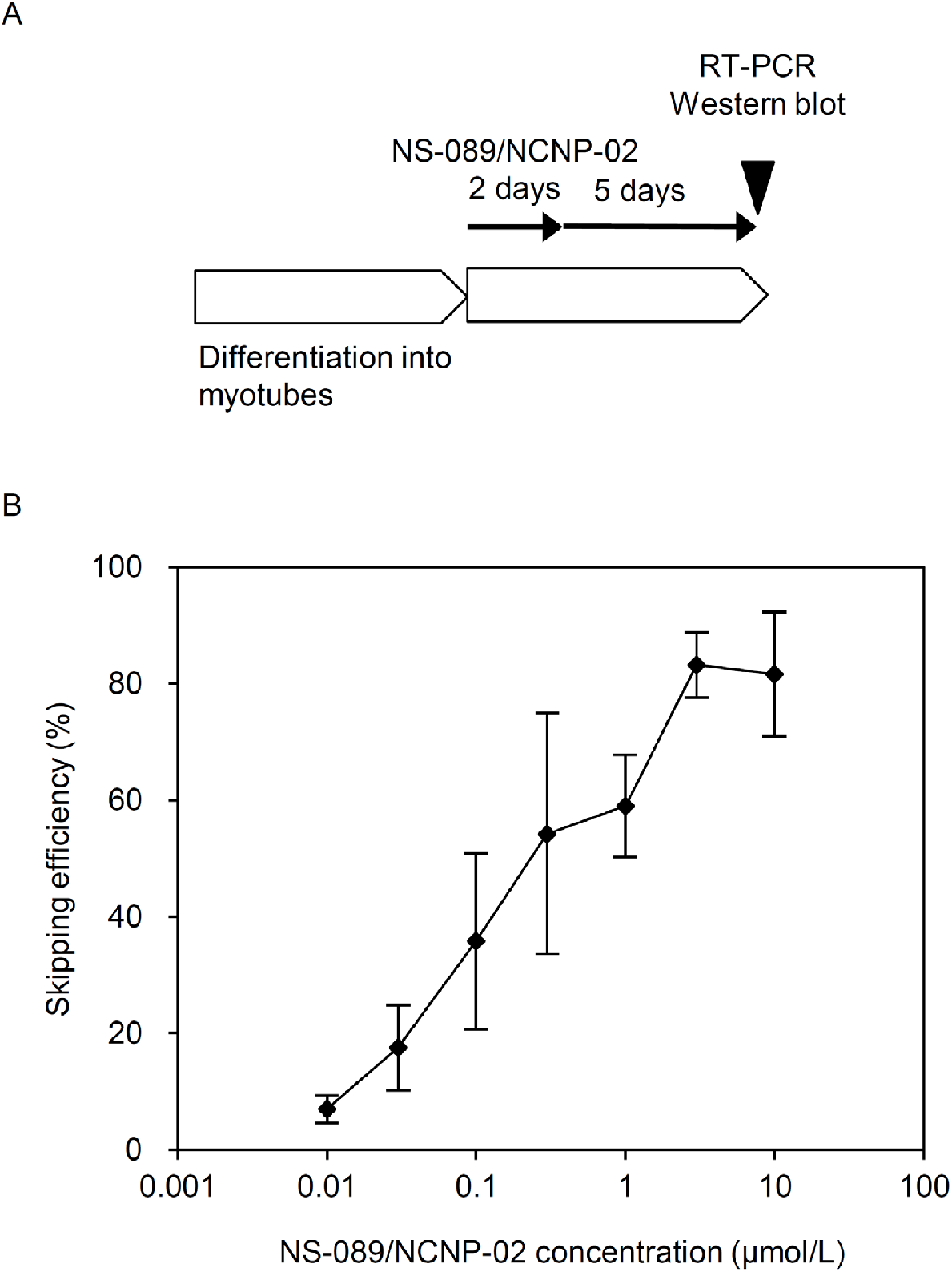
Exon 44 skipping activity of NS-089/NCNP-02 in cells derived from a DMD patient with a deletion of exon 45. (A) Schedule of PMO transfection for RT-PCR and western blotting. (B) Exon 44 skipping activity seven days after the start of a two-day treatment with NS-089/NCNP-02 of cells derived from a DMD patient with a deletion of exon 45. Each point shows the mean ± standard deviation (n = 4).

### NS-089/NCNP-02-induced expression of dystrophin protein

NS-089/NCNP-02-induced dystrophin protein expression was investigated in cells from a DMD patient with a deletion of exon 45. Patient fibroblasts were transduced with the human *MYOD* gene to induce differentiation into myotubes. The myotubes were then treated with NS-089/NCNP-02 at final concentrations of 0.01, 0.03, 0.1, 0.3, 1, 3 and 10 μmol/L for two days, after which the differentiation medium was replaced with medium containing no NS-089/NCNP-02. The expression of dystrophin protein was assessed by western blot analysis seven days after the beginning of treatment (Figure 3A). When NS-089/NCNP-02 was transfected into the cells at concentrations of 0.01, 0.03, 0.1, 0.3, 1, 3 and 10 μmol/L, the expression of dystrophin protein was 4.7%, 5.1%, 14.0%, 13.5%, 21.8%, 19.3% and 38.0%, respectively, of the expression of dystrophin protein in a cell lysate of myotubes differentiated from normal human fibroblasts by transduction with *MYOD*. (Figure 4A and 4B).

**Figure 4.**
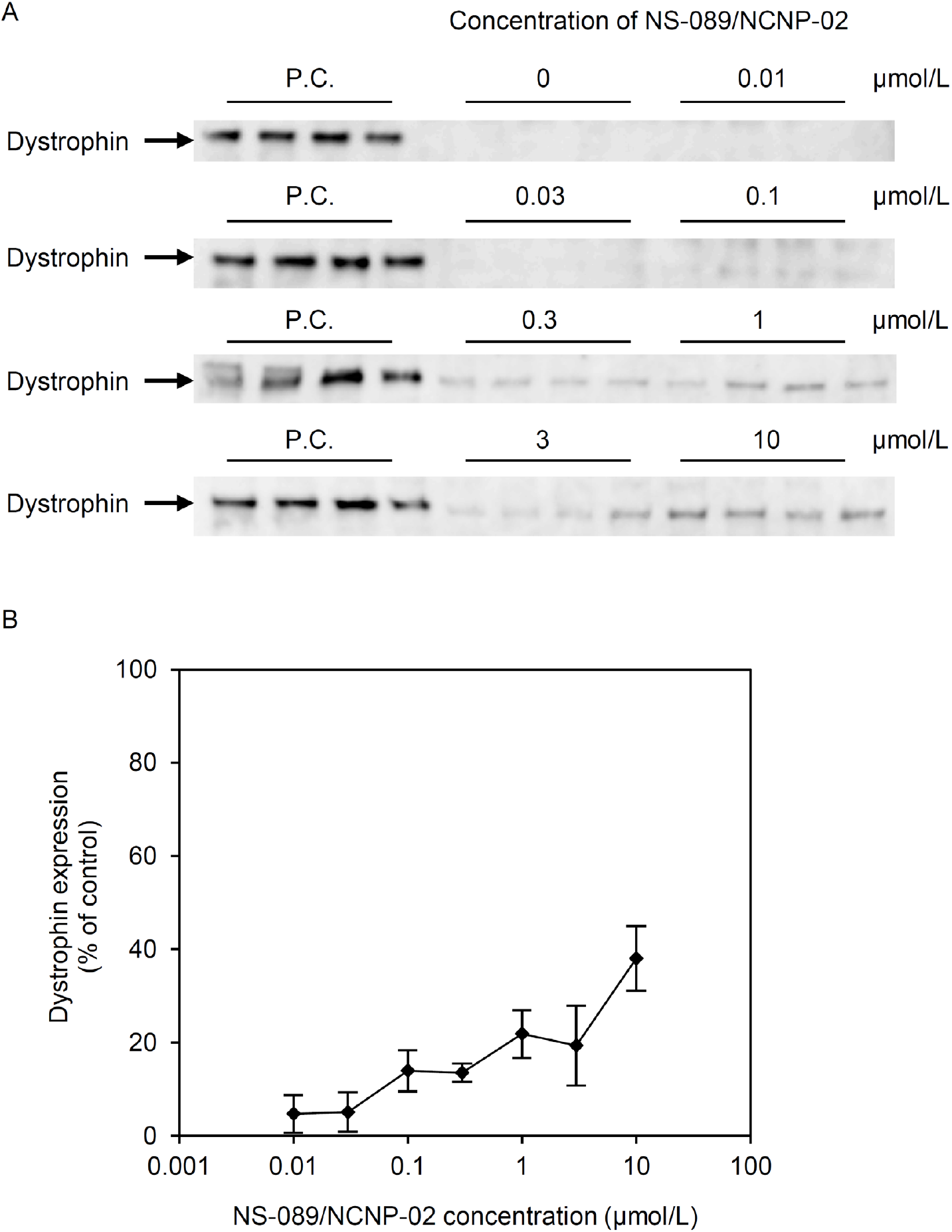
Expression of dystrophin protein seven days after the start of a two-day treatment with NS-089/NCNP-02 of cells derived from a DMD patient with deletion of exon 45. (A) Western blot analysis of dystrophin protein expression in the cells one week after transfection with NS-089/NCNP-02 over a period of two days. The positive control (P.C.) was a cell lysate of myotubes differentiated from normal human fibroblasts by transduction with *MYOD*. Samples were run in quadruplicate. (B) Densitometric analysis of the western blots relative to each P.C. on the same membrane. Each point shows the mean ± standard deviation (n = 4).

### Exon 44 skipping after transfection for a short incubation time

In the clinical setting, PMOs for DMD exon skipping are administered by weekly intravenous infusion. However, PMOs are known to be cleared from the blood with an elimination half-life of about 2 h after intravenous administration.^17^ To investigate exon 44 skipping activity after transfection for a short incubation time, the exon 44 skipping activity of NS-089/NCNP-02 after a 1-h transfection was measured in cells from a DMD patient with deletion of exon 45. Patient fibroblasts were transduced with the human *MYOD* gene to induce differentiation into myotubes. The myotubes were then treated with NS-089/NCNP-02 for 1 h at final concentrations of 0.01, 0.03, 0.1, 0.3, 1, 3 and 10 μmol/L. The exon 44 skipping activity was measured by RT-PCR seven days after the beginning of treatment (Figure 5A). NS-089/NCNP-02 induced exon 44 skipping with an EC_50_ value of 0.63 μmol/L (95% confidence interval, 0.40–0.98 μmol/L; Figure 5B).

**Figure 5.**
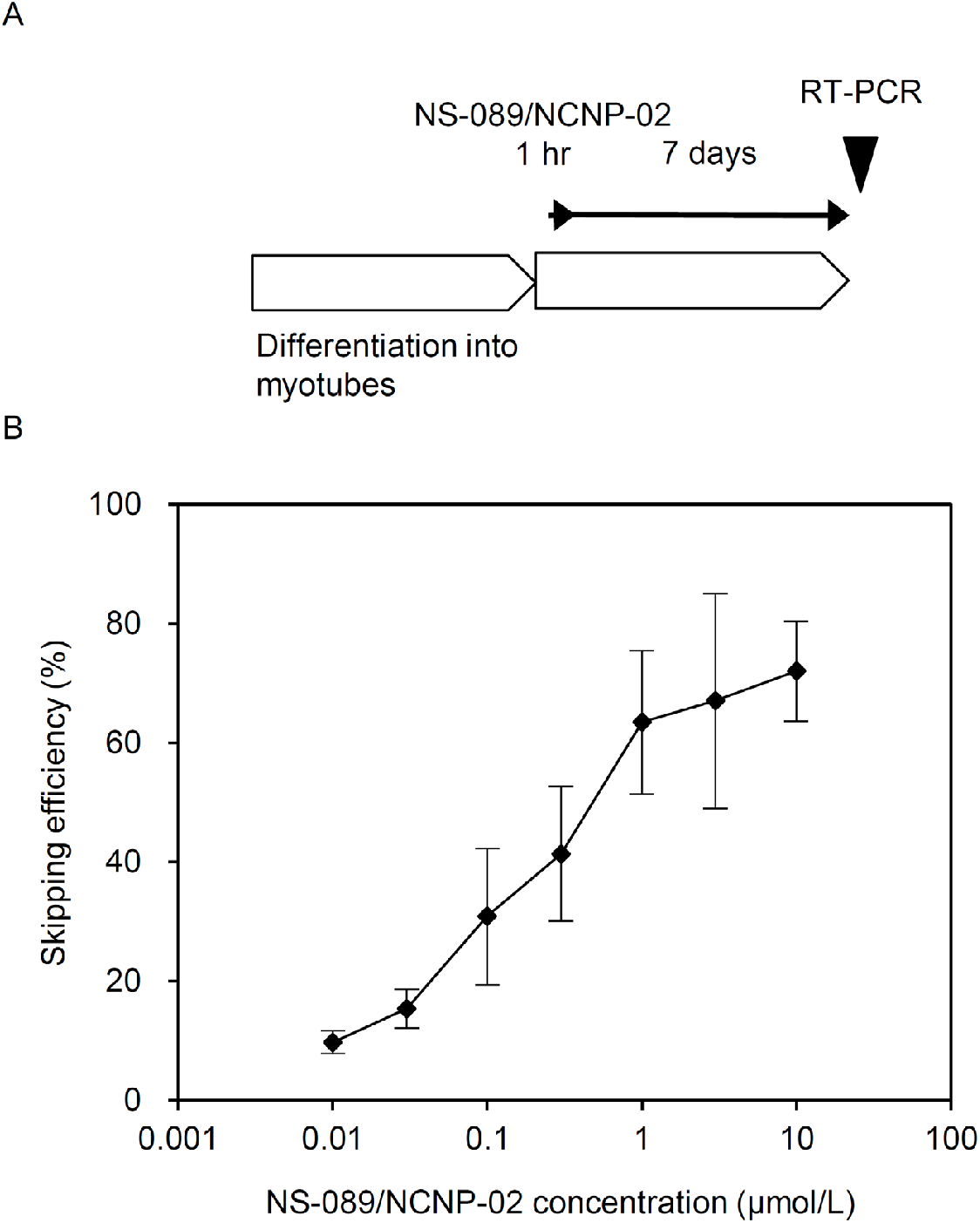
Exon 44 skipping activity seven days after the start of a 1-h treatment with NS-089/NCNP-02 of cells derived from a DMD patient with deletion of exon 45. (A) Schedule of PMO transfection for RT-PCR. (B) Each point shows the mean ± standard deviation (n = 4).

### Exon 44 skipping in cynomolgus monkeys

Finally, we assessed the *in vivo* activity of NS-089/NCNP-02 in cynomolgus monkeys. Although there is a single base difference between the target sequence of NS-089/NCNP-02 in cynomolgus monkeys and humans, we confirmed that NS-089/NCNP-02 showed exon 44 skipping activity in cultured monkey cells (data not shown). When NS-089/NCNP-02 was administered intravenously to cynomolgus monkeys once weekly for 13 weeks (Figure 6A), and the exon 44 skipping activity was measured in skeletal muscle (the right gastrocnemius muscle) the day after the last administration, the skipping efficiencies were 0.0%, 0.6%, 2.1% and 9.5% at doses of 0, 200, 600 and 2000 mg/kg, with significant skipping activity at 200, 600 and 2000 mg/kg (Figure 6B). The corresponding skipping efficiencies in cardiac muscle (the left ventricular apex) were 0.0%, 0.3%, 0.4% and 1.4%, with significant skipping activity at 600 and 2000 mg/kg (Figure 6C). In addition, NS-089/NCNP-02 was administered intravenously to cynomolgus monkeys once weekly for 13 weeks, and the exon 44 skipping activity was measured in skeletal muscle at the end of an eight-week recovery period (Figure 7A). The skipping efficiencies were 0.0%, 3.1% and 16.2% at doses of 0, 600 and 2000 mg/kg, with significant skipping activity at 2000 mg/kg (Figure 7B). The corresponding skipping efficiencies in cardiac muscle were 0.0%, 0.0% and 0.8%, and the skipping activity was not significant (Figure 7C).

**Figure 6.**
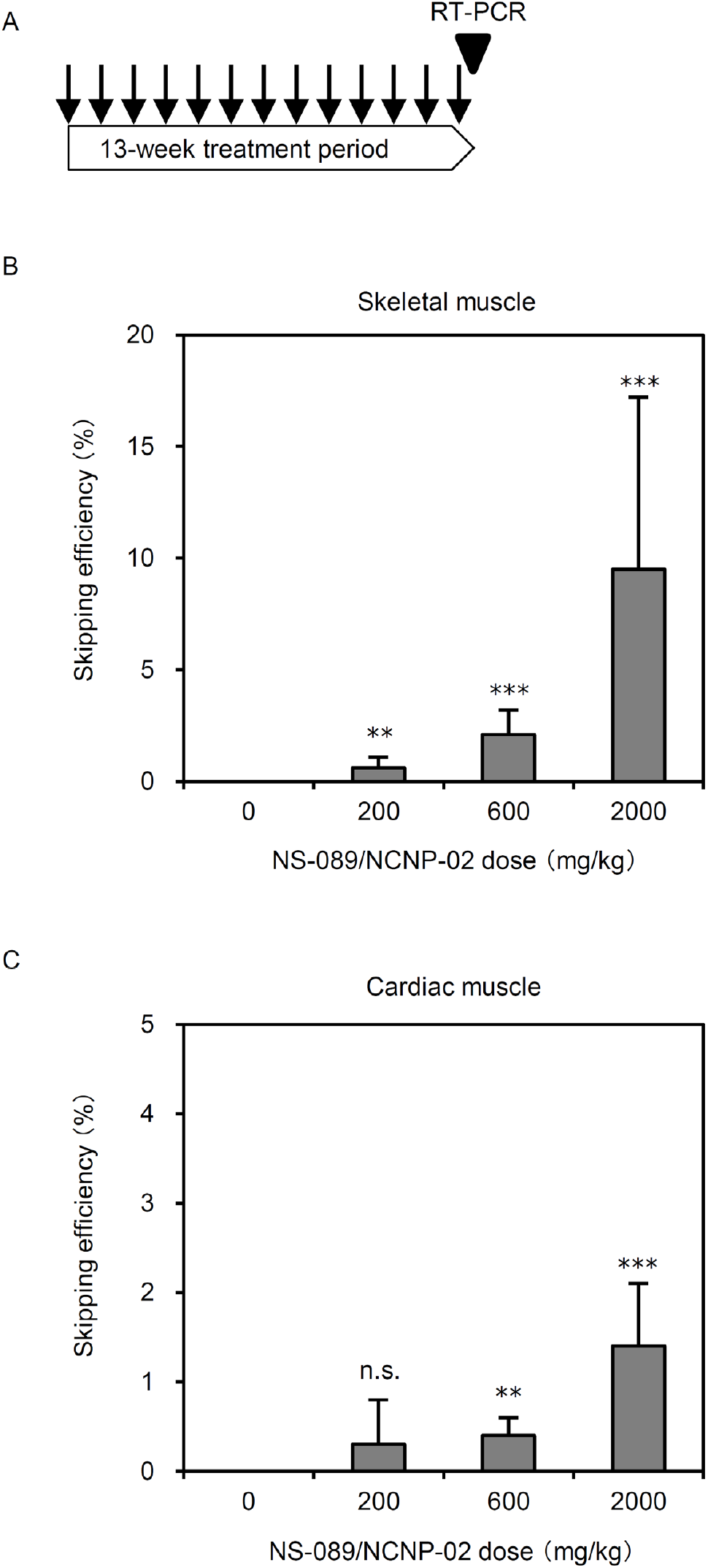
Exon 44 skipping activity in skeletal and cardiac muscle of cynomolgus monkeys at the end of a 13-week treatment period. (A) Schedule of PMO injection and RT-PCR. (B and C) Exon skipping was measured using RT-PCR in (B) skeletal muscle or (C) cardiac muscle. Each bar represents the mean and standard deviation (n = 5). Shirley–Williams multiple comparison test (one-sided) versus 0 mg/kg (saline) group. n.s., no significant difference; *, p < 0.005; ***, p < 0.0005.

**Figure 7.**
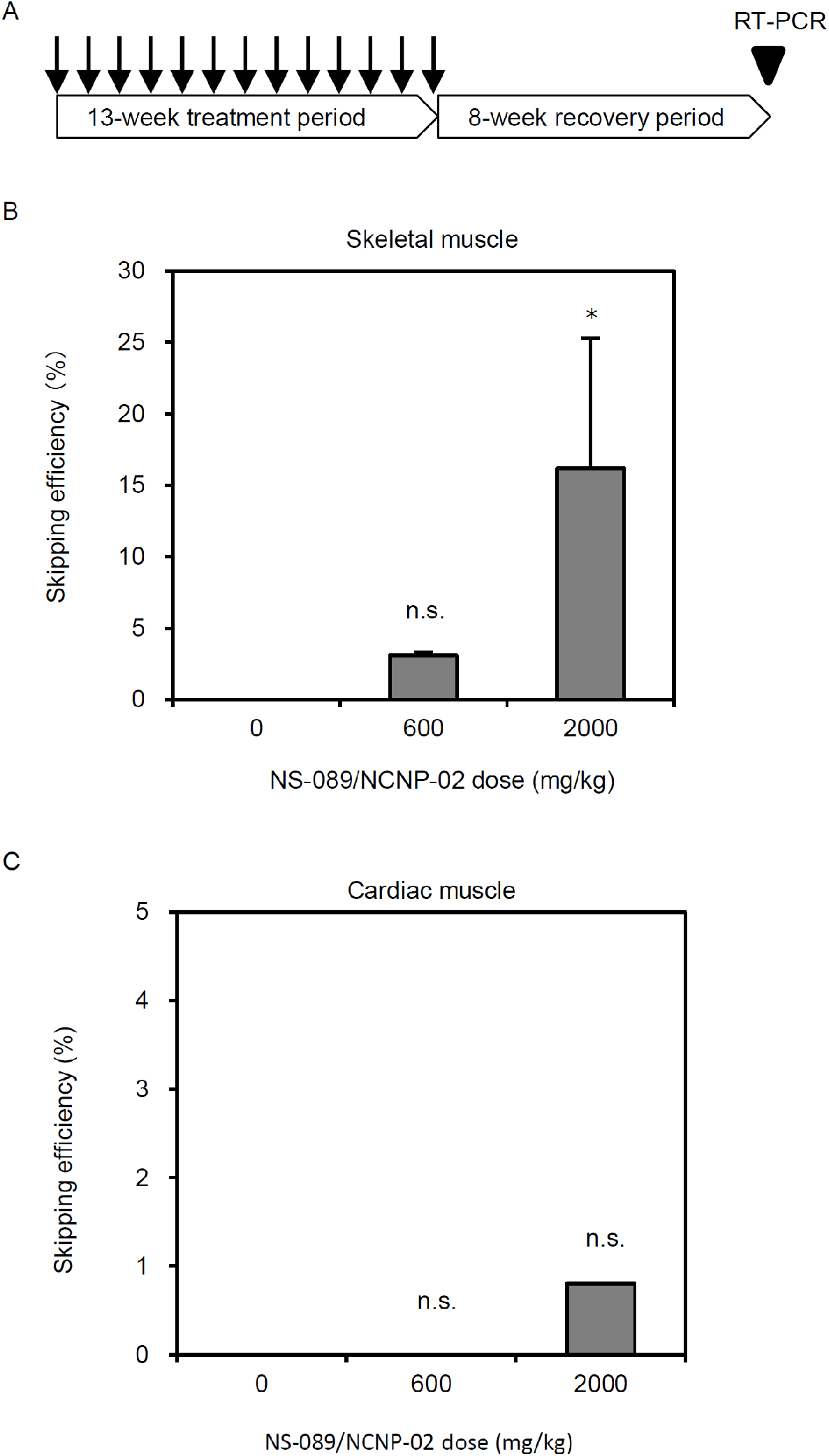
Exon 44 skipping activity in skeletal and cardiac muscle of cynomolgus monkeys at the end of an eight-week recovery period. (A) Schedule of PMO injection and RT-PCR. (B and C) Exon skipping was measured using RT-PCR in (B) skeletal muscle or (C) cardiac muscle. Each bar represents the mean and standard deviation (n = 2). Shirley–Williams multiple comparison test (one-sided) versus 0 mg/kg (saline) group. n.s., no significant difference.

## DISCUSSION

In this study, PMOs based on a novel design involving directly connected sequences targeting different parts of the exon were screened, and NS-089/NCNP-02 was found to have the highest exon 44 skipping activity. A combination of two ASOs has been reported to improve exon skipping activity.^15,16^ However, no previous report has shown that a single-strand ASO containing sequences targeting two or more sites in the same exon exhibits skipping activity. We have now found that enhanced exon 44 skipping activity was shown by 18–30-mer single-strand ASOs that target two separate sequences within exon 44. Positions 11–52 and 61–122 of exon 44 were identified as high skipping regions in the first screening. Combinations of positions 21–33 and 61–73 or 91–103 selected from positions 11–52 and 61–122, respectively, and a combination of positions 61–73 and 91–103, both selected from positions 61–122, showed high skipping activity. This novel type of ASO with two connected sequences expands the possibilities for creating highly active new sequences.

In contrast to the substantial exon 44 skipping activity shown by NS-089/NCNP-02, little activity was shown by a mixture of two separate 12-mer PMOs with exactly the same targeting sequences as NS-089/NCNP-02. Because PMOs less than 14 bases in length are expected to be inactive^18^, NS-089/NCNP-02 can be assumed to have induced exon 44 skipping by binding to both target sequences. The result obtained with NS-089/NCNP-02 is consistent with the idea that a single strand can simultaneously bind to two separate sites in the same exon and induce exon skipping.

NS-089/NCNP-02 showed exon 44 skipping activity and promoted dystrophin protein expression in *MYOD*-converted fibroblasts from a DMD patient. The EC_50_ value for the exon 44 skipping activity five days after a two-day treatment with NS-089/NCNP-02 was 0.33 μmol/L. The EC_50_ value for the exon 44 skipping activity seven days after a 1-h treatment with NS-089/NCNP-02 was 0.63 μmol/L. Thus, exon 44 skipping activity was sustained for seven days even when the contact time with the cells was shortened to 1 h. From these results, even though PMOs are cleared from the blood quickly after intravenous administration, NS-089/NCNP-02 can be expected to have an appreciable therapeutic effect. In the phase II study of viltolarsen in the US/Canada, dystrophin protein expression and exon 53 skipping activity were seen after treatment with viltolarsen, and preliminary results of timed function tests suggest clinical improvement in DMD boys.^19^ In a preclinical study, the EC_50_ value for the exon 53 skipping activity five days after a two-day treatment with viltolarsen delivered with the transfection reagent Endo-Porter was 0.82 μmol/L in cells from a DMD patient with deletion of exons 45–52.^20^ The EC_50_ value for the exon 44 skipping activity five days after a two-day treatment with NS-089/NCNP-02 without a transfection reagent was 0.33 μmol/L. The skipping efficiency was 31.9% one week after a 1-h treatment with viltolarsen at 10 μmol/L with Endo-Porter^20^ and 72.0% one week after a 1-h treatment with NS-089/NCNP-02 at the same concentration without a transfection reagent. Thus, NS-089/NCNP-02 showed higher exon 44 skipping activity than viltolarsen even though a transfection reagent was not used.

By exon 44 skipping, a dystrophin protein with the parts corresponding to exons 35–44, 44–45 or 44–54 deleted would be expected to be expressed in DMD patients with deletion of exons 35–43, 45 or 45–54, respectively.^21^ The shorter dystrophin protein induced by exon 44 skipping therapies is expected to be partially functional because deletion of exons 44–45 has only been reported three times, which suggests an ascertainment bias attributable to a very mild phenotype.^22^ Additionally, there are three reports of asymptomatic cases with deletions of exons 35–44, 38–44 and 41–44.^23–25^

To investigate the activity of NS-089/NCNP-02 *in vivo*, exon 44 skipping activity was measured in skeletal and cardiac muscle of cynomolgus monkeys. PMOs more efficiently enter fibers that are involved in active muscle regeneration, such as those which occur in DMD patients, than fibers of normal skeletal muscle.^26^ Even though exon 44 skipping is of low efficiency in the normal skeletal muscle of cynomolgus monkeys, exon 44 skipping is expected to be highly efficient in the skeletal muscle of DMD patients. Although it is generally difficult for PMOs to enter cardiomyocytes,^27,28^ some exon 44 skipping was induced by NS-089/NCNP-02 in the cardiac muscle of cynomolgus monkeys.

An investigator-initiated phase I/II study of NS-089/NCNP-02 demonstrated an increase in dystrophin protein expression and suggested maintenance of motor function or a trend in its improvement.^29^ Our data support these clinical results.

In conclusion, we describe the activity of NS-089/NCNP-02, a PMO for exon 44 skipping based on a novel design involving linked sequences targeting two sites in the same exon. NS-089/NCNP-02 induced exon 44 skipping and dystrophin protein expression both in cells derived from a patient with DMD amenable to exon 44 skipping and in the skeletal and cardiac muscle of cynomolgus monkeys.

## MATERIALS AND METHODS

### Antisense oligomers

2′-*O*-Methyl phosphorothioate oligonucleotides were purchased from Japan Bio Services Co., Ltd (Saitama, Japan). PMOs were synthesized by Nippon Shinyaku Co., Ltd.

### Cells

RD cells were obtained from the Health Science Research Resources Bank and cultured under 5% CO_2_ at 37°C in Eagle’s minimum essential medium (Sigma-Aldrich, St. Louis, MO, USA) containing 10% fetal bovine serum. Fibroblasts from a DMD patient with a deletion of exon 45 of the *DMD* gene (GM05112) were obtained from the Coriell Institute for Medical Research. Fibroblasts were cultured and induced to differentiate into myotubes by *MYOD* conversion as previously described.^20^

### Transfection of PMOs into cells

PMOs were dissolved in distilled water and transfected into RD cells using the Amaxa Cell Line Nucleofector Kit L and a Nucleofector II electroporation device (Lonza, Basel, Switzerland) with program T-030 or into cells from a DMD patient without a transfection reagent.

### Reverse transcriptase polymerase chain reaction (RT-PCR)

Total RNA was extracted from RD cells or cells from a DMD patient, and RT-PCR was performed as previously described.^18^ From approximately 15-mg pieces of tissue obtained from 2-year-old male cynomolgus monkeys in a 13-week intermittent intravenous dose toxicity study of NS-089/NCNP-02 followed by an 8-week recovery period (study approved by the Institutional Animal Care and Use Committee of Nippon Shinyaku Co., Ltd; approval no. 17062701), total RNA was extracted using TissueLyser II (Qiagen, Valencia, CA, USA) and NucleoSpin RNA (Macherey-Nagel, Düren, Germany). RNA concentrations were determined from the absorbance at 260 nm using a NanoDrop ND-1000 spectrophotometer (Thermo Fisher Scientific, Waltham, MA, USA). RT-PCR was performed with 200 ng of extracted total RNA using a Qiagen OneStep RT-PCR Kit (Qiagen). The primers used were a forward primer (Hokkaido System Science, Sapporo, Japan) designed for exon 43 (5’-GCTCAGGTCGGATTGACATT-3’) and a reverse primer (Hokkaido System Science) designed for exon 47 (5’-GGGCAACTCTTCCACCAGTA-3’) so as to exclude exon 44. RT-PCR was performed with a Takara Thermal Cycler Dice^®^ Touch (Takara Bio, Kusatsu, Japan). The RT-PCR program was as follows: reverse transcription at 50°C for 30 min, heat denaturation at 95°C for 15 min and 35 cycles consisting of denaturation at 94°C for 1 min, annealing at 60°C for 1 min, and extension at 72°C for 1 min, followed by a final extension at 72°C for 10 min. The PCR reaction products were analyzed using a 2100 Bioanalyzer (Agilent Technologies, Waldbronn, Germany). The skipping efficiency was determined from the molarity of the PCR products by the following expression: (PCR reaction products without exon 44) × 100 / [(PCR reaction products without exon 44) + (PCR reaction products with exon 44)].

### Western blotting

Western blotting was performed on lysates of cells from a DMD patient, and the results were analyzed as previously described.^20^

### Statistical analysis

All analyses were performed using SAS software (Ver. 9.3; SAS Institute, Cary, NC, USA) and EXSUS (Ver. 8.0.0; CAC Exicare, Tokyo, Japan). The skipping efficiencies obtained were analyzed by nonlinear regression using a two-parameter logistic model to calculate EC_50_ values.

## Author contributions

N.W., Y.T., T.N., S.M. and T.S. performed the experiments, N.M., T.N., K.T., Y.A. and S.T. coordinated and supervised the project, and N.W. wrote the manuscript.

## Data availability

All data are included in the manuscript. Raw data are available on request.

## Acknowledgments

The authors wish to thank Dr. Gerald E. Smyth for his assistance in preparing the manuscript. The work was performed in Kodaira, Tokyo, and Tsukuba, Ibaraki, Japan.

---

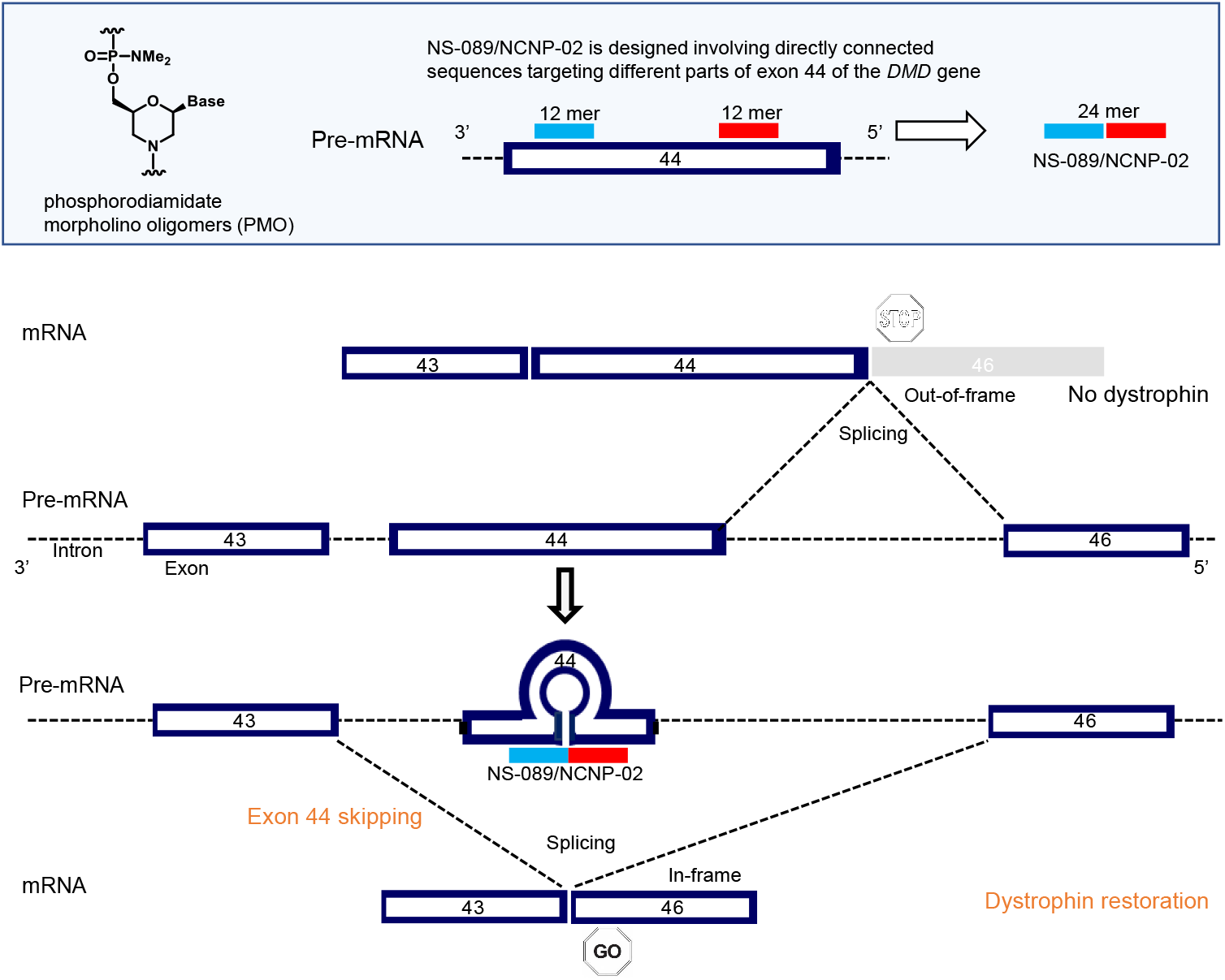

